# Mutational signatures in countries affected by SARS-CoV-2: Implications in host-pathogen interactome

**DOI:** 10.1101/2020.09.17.301614

**Authors:** J. Singh, H. Singh, S E. Hasnain, S.A. Rahman

## Abstract

We are in the midst of the third severe coronavirus outbreak caused by SARS-CoV-2 with unprecedented health and socio-economic consequences due to the COVID-19. Globally, the major thrust of scientific efforts has shifted to the design of potent vaccine and anti-viral candidates. Earlier genome analyses have shown global dominance of some mutations purportedly indicative of similar infectivity and transmissibility of SARS-CoV-2 worldwide. Using high-quality large dataset of 25k whole-genome sequences, we show emergence of new cluster of mutations as result of geographic evolution of SARS-CoV-2 in local population (≥10%) of different nations. Using statistical analysis, we observe that these mutations have either significantly co-occurred in globally dominant strains or have shown mutual exclusivity in other cases. These mutations potentially modulate structural stability of proteins, some of which forms part of SARS-CoV-2-human interactome. The high confidence druggable host proteins are also up-regulated during SARS-CoV-2 infection. Mutations occurring in potential hot-spot regions within likely T-cell and B-cell epitopes or in proteins as part of host-viral interactome, could hamper vaccine or drug efficacy in local population. Overall, our study provides comprehensive view of emerging geo-clonal mutations which would aid researchers to understand and develop effective countermeasures in the current crisis.

**Significance:** Our comparative analysis of globally dominant mutations and region-specific mutations in 25k SARS-CoV-2 genomes elucidates its geo-clonal evolution. We observe locally dominant mutations (co-occurring or mutually exclusive) in nations with contrasting COVID-19 mortalities per million of population) besides globally dominant ones namely, P314L (ORF1b) and D164G (S) type. We also see exclusive dominant mutations such as in Brazil (I33T in ORF6 and I292T in N protein), England (G251V in ORF3a), India (T2016K and L3606F in ORF1a) and in Spain (L84S in ORF8). The emergence of these local mutations in ORFs within SARS-CoV-2 genome could have interventional implications and also points towards their potential in modulating infectivity of SARS-CoV-2 in regional population.

## Introduction

The rapid spread of SARS-CoV-2 has created an unprecedented global crisis. The evolving SARS-CoV-2 viral variants have shown dominance (present in >50% genomes) of some mutations indicating their concurrent emergence worldwide. However, co-dominance of geographically distinct mutations may hamper global vaccine development and pharmacological interventions. The genome organization of the 29kb SARS-CoV-2 (1) involves 12 Open Reading Frames (ORF) encoding for ORF1a and 1b, surface glycoprotein - Spike (S), ORF3a, envelope (E), membrane (M), ORF6, ORF7a and 7b, ORF8, nucleocapsid (N) proteins and ORF10 (2,3). Our earlier phylogenetic analysis of >250 SARS-CoV-2 isolates showed clustering of clinical samples in multiple clades derived from the molecular divergence in ORF1ab/1b, S and ORF8 proteins (4). Subsequent analysis of isolates sequenced in India showed higher mutational frequency in ORF1a from Indian samples (n=25) compared to global isolates (n=3932) (5). This prompted us to look for similar dominant mutations, concurrent or unique, present both globally and unique to nations with higher COVID-19 incidence namely Brazil, India, Italy, Spain, UK and USA and compare it to countries where incidence was low, Australia, Germany, Republic of Congo and Saudi Arabia. Here, we provide a comprehensive report on the evolution of local clusters of mutations explored in a large dataset of >25k genomes (whole genome sequences) from multiple geographical locations around the globe and their co-occurrence or exclusivity with common global mutations. We further ascertained effects of these mutations on the structural stability in the corresponding proteins, some of which interact with host proteins, which were shown to be upregulated in patients with severe COVD-19 symptoms. Identification of such mutations may further aid researchers in developing broad spectrum vaccines or understanding variability in drug efficacy.

## Methods

High quality, full length whole genome sequences of clinical isolates of SARS-CoV-2 were downloaded from Global Initiative on Sharing All Influenza Data (Table 1) (GISAID; https://www.gisaid.org) (23). Glimmer was used for gene prediction (ORFs), and ORF sequences with Ns were masked out and were not considered for protein translation (24). Throughout, site numbering and genome structure are given using Wuhan-Hu-1 (NC_045512.2) as reference genome (https://www.ncbi.nlm.nih.gov/nuccore/NC_045512).

**Table 1:**
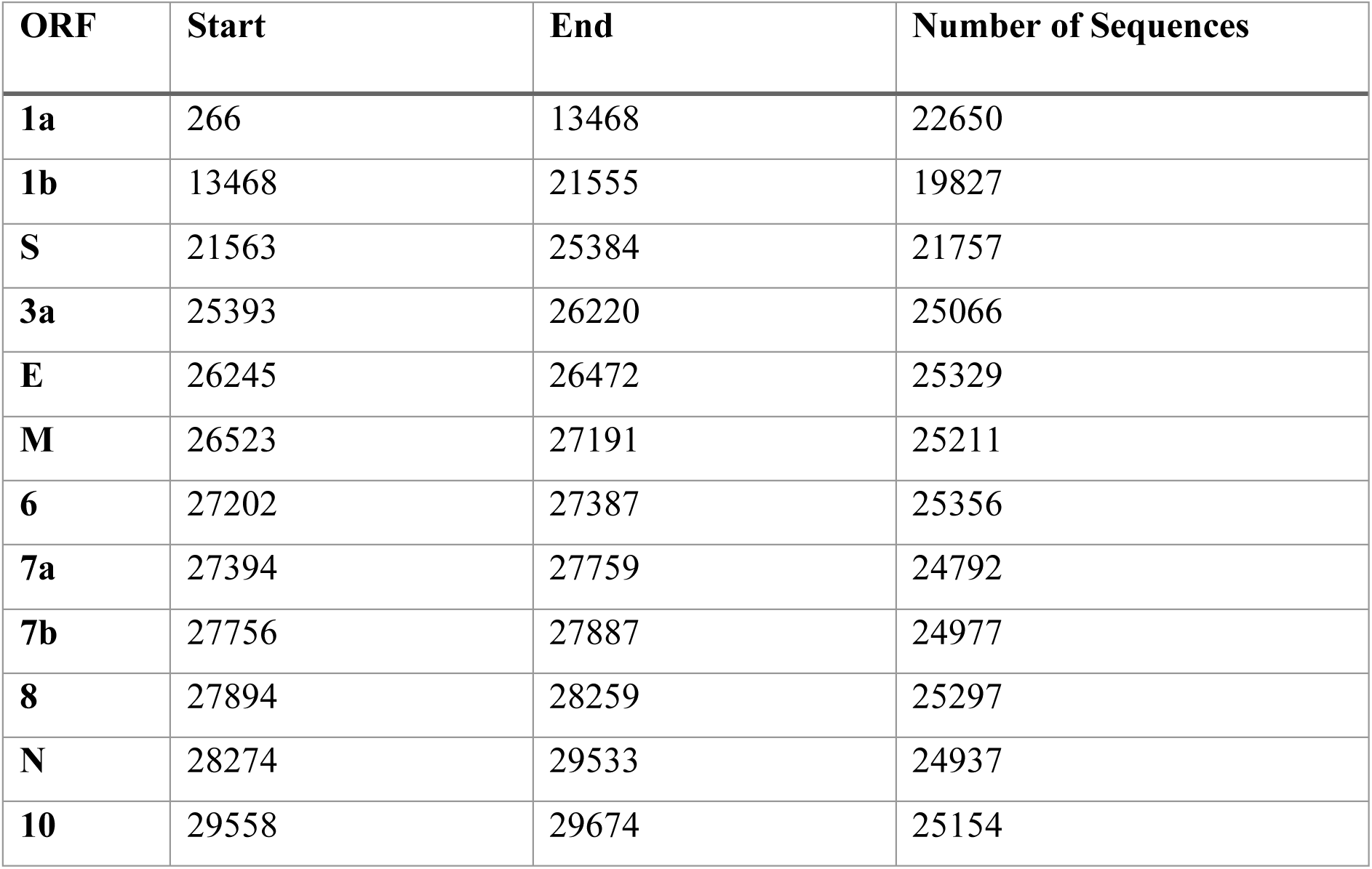
Distribution of ORFs (amino acid sequences) in SARS-CoV-2 genomes used in this study. The start and end positions for respective ORFs were taken on the basis of gene coordinates of reference genome NC_045512.

| ORF | Start | End | Number of Sequences |
| --- | --- | --- | --- |
| **1a** | 266 | 13468 | 22650 |
| **1b** | 13468 | 21555 | 19827 |
| **S** | 21563 | 25384 | 21757 |
| **3a** | 25393 | 26220 | 25066 |
| **E** | 26245 | 26472 | 25329 |
| **M** | 26523 | 27191 | 25211 |
| **6** | 27202 | 27387 | 25356 |
| **7a** | 27394 | 27759 | 24792 |
| **7b** | 27756 | 27887 | 24977 |
| **8** | 27894 | 28259 | 25297 |
| **N** | 28274 | 29533 | 24937 |
| **10** | 29558 | 29674 | 25154 |

MAFFT was used for ORF alignment and mutations were computed using BioInception’s in-house pipeline based on R (https://cran.r-project.org/) and Python. Alignment regions with >25% gaps against the reference genome were ruled out for amino acid mutational changes. The change or mutation ratio for an ORF alignment is computed by the presence of a mutation divided by total number of sequences in the alignment. This ratio is changed into a percentage. Analysis of co-occurring or exclusive mutations (number of times two mutation occur together in a clinical genome) across various ORFs was defined in terms of log-odds ratio (with p-value and standard error) where a positive ratio indicates co-occurring mutations and negative value denotes an exclusive mutation in the significant part of population using statsmodel (Supplementary Data Sheet 1) (25). Structural effects of mutations on protein conformation, flexibility and stability were predicted using DynaMut (26). The structural models were retrieved from i-tasser COVID-19 database (27).

## Results

### Mutations occurring globally in ≥25% of the population

Here, we report recurrent occurrence of (i) global and (ii) local (region specific) non-synonymous mutations present in >25K SARS-CoV-2 clinical isolates (Table 1). To achieve better signal-to-noise ratio, we have used a double layered protocol to identify statistically significant signature mutations. In general, recurrent or dominant mutations occurring in ≥10% of the functional genome (ORFs), compared to Wuhan_HU-1 (NC_045512), were classified into mutually exclusive or co-occurring (mutations which also have the propensity to exist together with another mutation). These were defined using log-odds ratio (LOR) where a positive or negative log-odds value represents co-occurring and mutually exclusive mutations, respectively. Vibrational entropy changes were predicted for both wild type and mutant proteins for evaluating the impact of mutations on conformation, flexibility and stability of proteins. The SARS-CoV-2 proteome can be classified into ORF1a, ORF1b (or ORF1ab joint), S, E, ORF7a, ORF7b, N, ORF8 and ORF 10. ORF1a and ORF1b code for polyproteins which are further cleaved to non-structural proteins (nsp), necessary for infection and replication of virus. Our analysis revealed high propensity mutations present in the ORF1a and 1b, S, ORF3a, ORF6, ORF8 and N proteins of SARS-CoV-2 (Figure 1, Supplementary Table 1). Our findings agree with earlier reports on global co-occurrence of P314L (70.97%) and D614G (71.57%) mutations in ORF1b (corresponding to P323L in nsp12) and S proteins, respectively.

**Figure 1.**
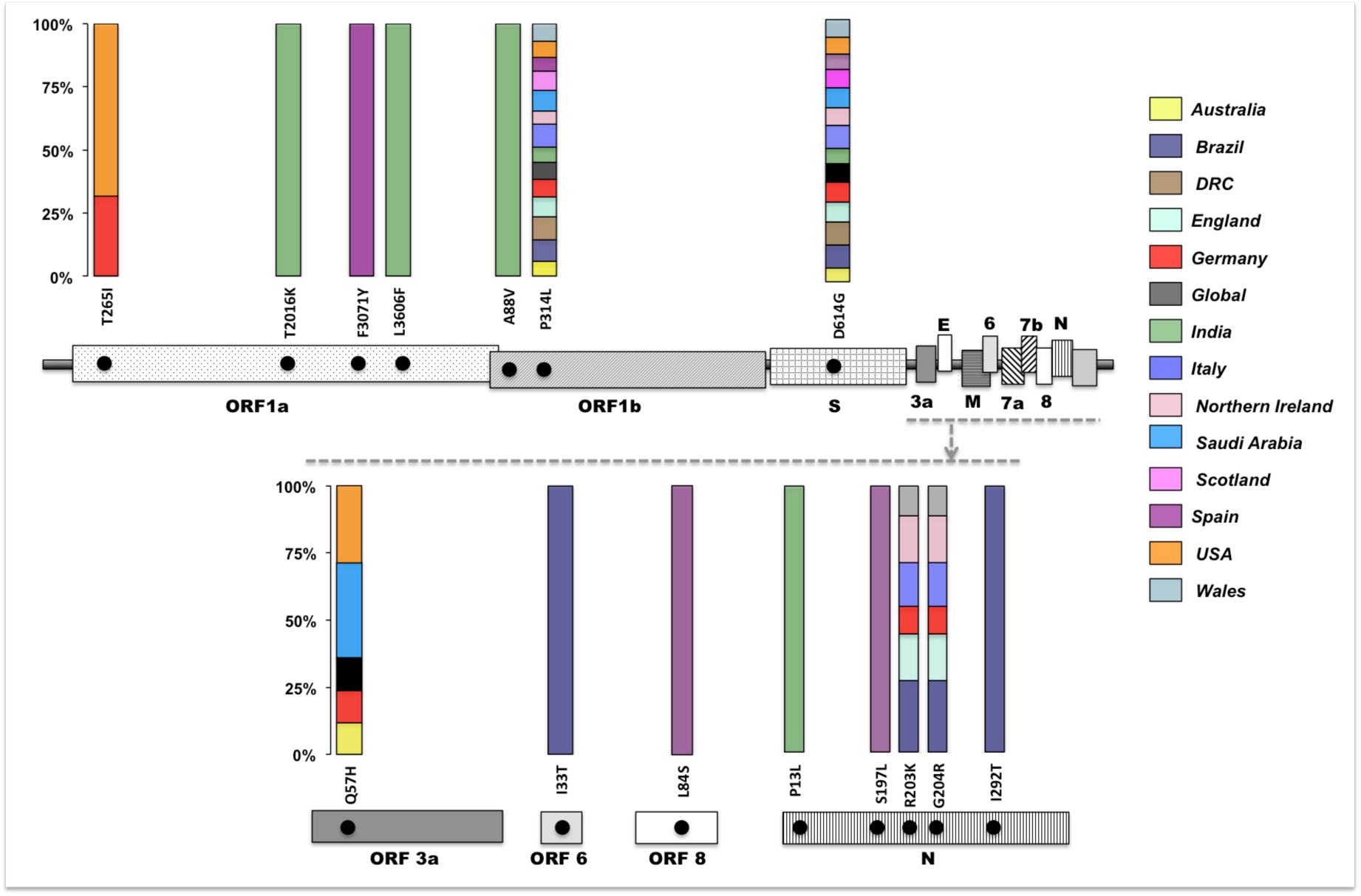
Mutation analysis of ORF’s predicted for 25K SARS-CoV-2 genomes. Position of mutations present in ≥25% of genome samples in different ORF’s along with stacked percent occupancy of these mutations. Single stacked occupancy highlights mutations specifically observed in samples from respective nations. e.g. T2016K, L3606F (ORF1a) and P13L in N protein are specifically observed in the Indian Isolates while F3071Y (ORF1a) and L84S (ORF8) are observed in Spain.

Subsequently, high occurrence of mutation Q57H (25.54%) in ORF3a protein, R203K (24.34%) and G204R (24.29%) in N protein (Supplementary Table 1) was globally reported. They were also found to be widely accumulated in North American and European populations (6).

### A comparative study between region specific mutations and globally dominant mutations

By capturing the most prominent mutations present in samples (through application of a coarse data filter for mutations present in >25% of samples), our hierarchical (ward based) clustering analysis of regional isolates placed European nations (except Spain and Scotland) and Brazil into one cluster and Australia, India, Saudi Arabia, Scotland, Spain, USA, and DRC into another cluster (Figure 2). Brazil and India, however, do not cluster with the rest of populations and seem to possess distinct signature mutations, indicating early signs of genetic drift. In isolates obtained from populations with the incidence of over a hundred mortalities per million e.g. Italy, England, Wales, we observed positive log-odds ratio amongst mutations P314L-D614G (LOR: 10.75, p-value 0.0) and R203K-G204R (LOR: 14.39, p-value 4.089e-146). Interestingly Germany and Italy (with contrasting mortalities per million of population) (7) show similar mutation signatures in their genomes. However, in relatively high viral burden populations such as in Spain, we observed mutually exclusive presence of L84S (ORF8) mutation along with globally occurring mutations P314L (ORF1b) and D614G (S) (Figure 1). Whereas in the USA, T265I (ORF1a) and Q57H (ORF3a) were highly concurrent with P314L-D614G mutations. The globally dominant P314L, D614G and Q57H were predicted to be stabilizing mutations for nsp12, S and ORF3a proteins, respectively (Table 2). Similarly, other mutations were predicted to confer mild stabilization or destabilization effects on respective proteins (ΔΔG -0.64 to 0.46 kcal.mol^-^1) (Table 2).

**Table 2:** Predictions on effect of mutations on protein stability and flexibility. Positive and negative ΔΔG indicate an overall stabilization or destabilization effect of mutation on tertiary architecture, respectively. ΔΔG_vib_ (ENCoM) indicates vibrational entropy energy between wild type and mutant and dictates effects of mutation on protein flexibility. Positive and negative ΔΔG_vib_ indicate increase and decrease in molecule flexibility, respectively.

| ORF | Mutation | $\Delta\Delta G$<br>kcal.mol<sup>-1</sup> | $\Delta\Delta G_{\text{vib}}$ (ENCoM)<br>kcal.mol<sup>-1</sup>.K<sup>-1</sup> | Mutation Effect |
| --- | --- | --- | --- | --- |
| **1a** | T265I | 0.09 | 0.02 | Stabilizing |
|  | T2016K | -0.64 | 0.43 | Destabilizing |
|  | L3606F | 0.35 | -0.18 | Stabilizing |
| **1b** | P314L | 0.46 | -0.32 | Stabilizing |
| **3a** | Q57H | 0.07 | -0.40 | Stabilizing |
| **S** | D614G | 0.25 | 0.20 | Stabilizing |
| **6** | I33T | -0.04 | 0.58 | Destabilizing |
| **8** | L84S | -0.345 | 0.03 | Destabilising |
| **N** | P13L | 2.25 | -1.16 | Stabilizing |
|  | S197L | 0.382 | 0.05 | Stabilizing |
|  | R203K | -0.25 | 0.47 | Destabilizing |
|  | G204R | 0.23 | -1.19 | Stabilizing |
|  | I292T | -1.99 | 0.77 | Destabilizing |

**Figure 2.**
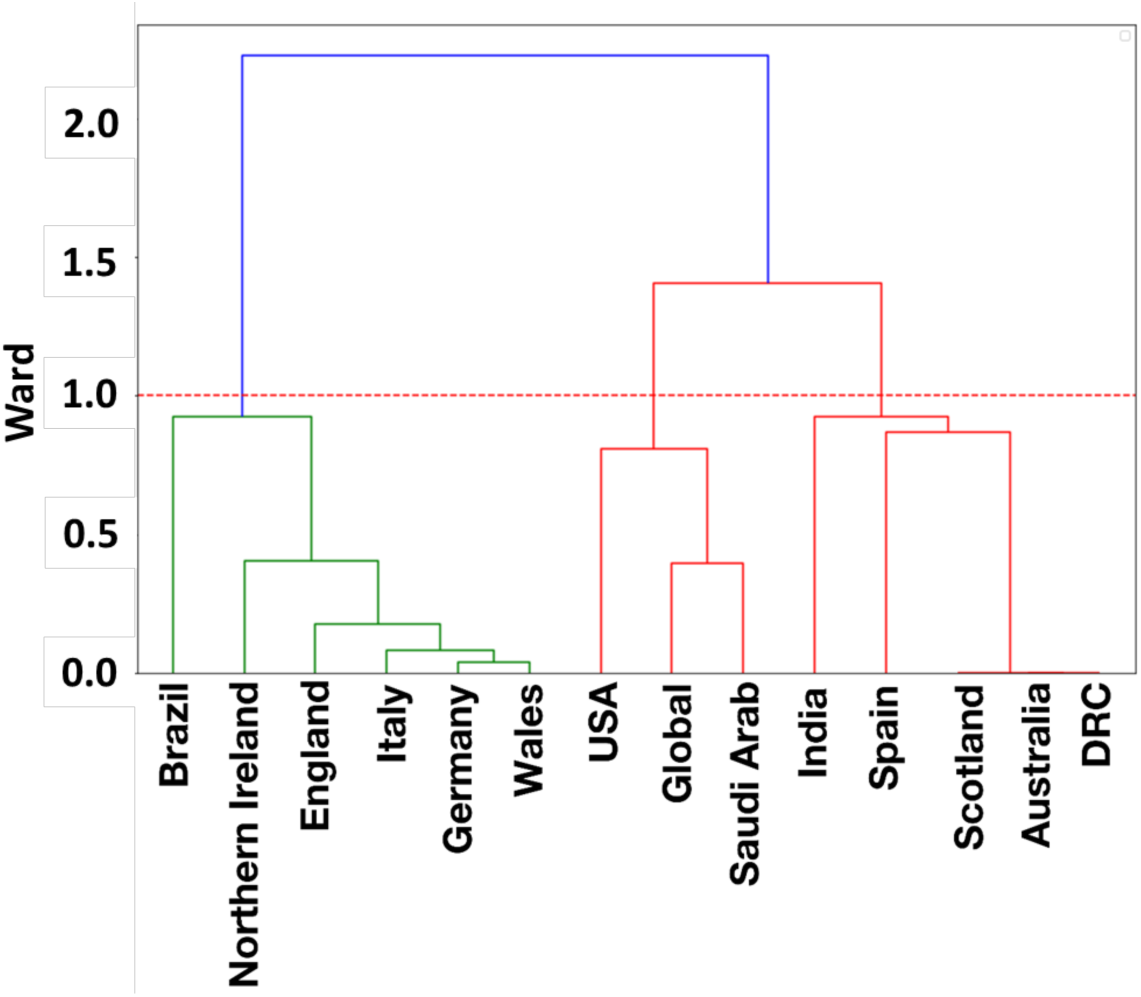
Hierarchical (Ward) Clustering of regional isolates based on propensity (present in ≥25% of samples) of mutations to co-occur or be mutually exclusive. Two major groups can be observed based on co-occurring and mutually exclusive mutations. India and Brazil segregate from both groups owing to some country specific mutations.

Specific (present only in the particular region within the cut-off limit) mutations - L3606F (corresponding to L37F in nsp6) (30.96%) and T2016K (corresponding to T1198K in nsp3) (26.61%) both in ORF1a, A88V (corresponding to A97V in nsp6) (25.67%) in ORF1b and P13L (31.24%) in Nucleoprotein (N protein) were present in Indian isolates and did not co-occur with P314L, D614G type (Supplementary Table 1, Supplementary Data Sheet 1). The outlier Indian samples could be attributed to these mutations which can account for an entirely separate Indian clade. Though the L3606F mutation also occurs in 25% of isolates from Australia, DRC, England and Wales, the P13L, T2016K mutant strains seem to have newly co-evolved with L3606F variants in later stages of transmission in India. Brazil, another outlier, has two distinct mutations I33T (52.53%) in ORF6 and I292T (59.6%) in N protein which were absent in other nations. These co-occurred with P314L and D614G unlike the Indian exclusive mutations L3606F, T2016K and P13L which did not co-occur. The Brazilian exclusive mutations I33T and I292T co-occurred with P314L, D614G, R203K and G204R. It is also worth noting that within the N protein, the Indian and Brazilian isolates show different country specific mutations - P13L and I292T mutations, respectively. Stability analysis on these mutations predicted induction of significant and contrasting conformational changes in N protein. The P13L mutation was predicted to induce high stabilization (ΔΔG 2.25 kcal.mol^-1^) owing to hydrophobic interactions of mutated Leu with Ala152 (Supplementary Figure 1). On the contrary, the I292T (exclusive in Brazil) was predicted to destabilize N protein (ΔΔG -1.99 kcal.mol^-1^). The destabilization was observed due to changes in weak H-bond interactions in the vicinity of T292 (Supplementary Figure 2).

### Mutations occurring in more than 10% of population

While prominent mutations (by applying a coarse data filter for mutations present in >25% of samples) yielded specific mutations in isolates from India, Brazil, USA and Spain, further reducing the cut-off to ≥10% showed additional country specific signature mutations in different ORFs (Figure 3). Geographically defined genomic evolution is expected. In United Kingdom, country specific mutations were observed in England, Wales, Northern Ireland and Scotland. Mutations in England (G251V in ORF3a), Wales (G392D, A876T in ORF1a; S193I in N protein), Scotland (S194L in N protein) and Northern Ireland (P2144L and T3579I in ORF1a) displayed mutual exclusivity (negative LOR) with P314L, D614G and R203K, G204R mutations. Interestingly in England, the G251V mutation was highly concurrent with L3606F mutation of nsp6. Contrarily in Wales, the G392D, A876T and S193I mutations (which have high propensity to co-occur together, showed mutual exclusivity with the L3606F mutation (Supplementary Data Sheet 1). The P2144L and T3579I mutations showed a high propensity to co-occur in Northern Ireland population. Distribution of co-occurring and exclusive mutations in United Kingdom clearly shows G392D, A876T, S193I for Wales and P2144L, T3579I as specific for Northern Ireland, distinct from England.

**Figure 3.**
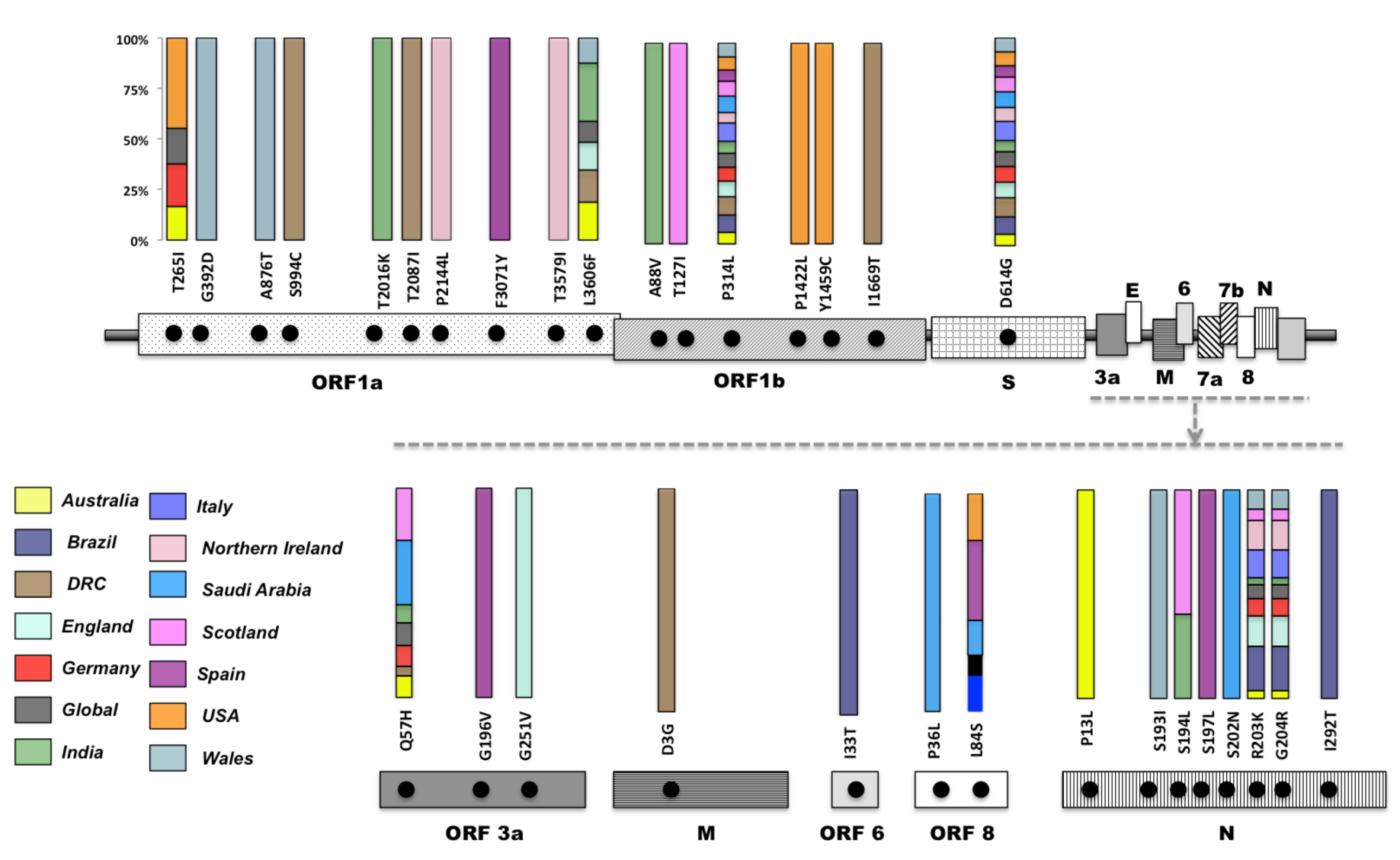
Position of mutations present in ≥10 % (including the >25% mutations as depicted in Figure 1a) of genome samples in different ORF’s along with stacked percent occupancy of these mutations. Single stacked occupancy highlights mutations specifically observed in samples from respective nations.

Antigenic drift is believed to occur due to immune pressure resulting in selection of viruses that can escape, hence exploring mutations will aid towards vaccine design and development (8). Virus circulating in humans undergo antigenic drift, thus necessitating periodic updates to the virus strains contained in the seasonal vaccine to maintain a good match with circulating viruses (9) (10). In general, variations in the virus can lead to reduced efficacy and narrow protection from most vaccinations. The mutating strains might weaken the memory B cells that have the capacity to generate and improve antibody response to new infection. A correlation of the occurrence of hotspot mutations within potential T cell and B cell epitopes revealed that F3071Y and L3606F (ORF1a) are within the epitope regions predicted for both CD4+ and CD8+ T cell activity (11). The R203K and G204R from N protein have a strong propensity for B cell epitopes.

## Discussion

The viral mutation rates (12) are important for understanding the tractability of using variable regions as vaccine candidates and how virus populations will respond to the application of a mutagen (13). Bioinformatics and computational biology can assist in the discovery of conserved epitopes through sequence variability analysis (14). This is particularly relevant when dealing with virus capable of evading the immune system due to their high mutation rates. These approaches, combined with experimental evolution and deeper mechanistic studies, will aid in the understanding of whether mutation rates are likely to change in the future. While our observations on geographical distribution of mutations could indicate significant public health outcomes, these also raises fundamental questions on the likely basis for emergence of such mutations as a function of geography. A number of factors such as differences in the microbiome population in a given region, drug pressure, and host genetics could be attributed. In separate studies on various human infecting viruses, it has been shown that commensal microbiome can prime the viral infections in hosts (15). The ongoing long and short term evolutions in the microbiome may further add complexity to our understanding of host shaped viral adaptations (16). Drug selection pressures can also work at regional and sub-regional levels (17,18). Analysis of mutation rates are equally crucial in terms of protein-protein interactions within coronavirus ORFeome and host-pathogen interaction interface. In SARS-CoV, nsp2, nsp8 (both part of ORF1ab) and ORF9b were shown to interact with other viral proteins (19). For instance, nsp8 (part of ORF1ab) showed prominent interactions with replicase proteins including nsp2, nsp5, nsp6, nsp7, nsp9, nsp12, nsp13 and nsp14 (all part of ORF1ab), indicating crucial role of intra-viral protein-protein interactions in replication machinery. The mutations in ORFs could be equally crucial in terms of host-viral interactome. Shen Bo *et. al*. (20) classified severity of COVID-19 on the basis of differential expression of 93 serum proteins and 204 metabolite signatures, highlighting immune and metabolic dysregulation in COVID-19 patients.

Another study by Gordon *et. al*. (21) identified physical interactions of host (human) proteins with SARS-CoV-2. Protein-protein interactome (PPI) analysis yielded 332 high confidence druggable human proteins which are involved in protein trafficking, transcription-translation and ubiquitination regulation. We observed three common proteins in these studies (20,21) which are both high confidence druggable targets and dysregulated between healthy and severe COVID-19 symptoms. Two of these proteins; CD antigen CD155 and gamma-Glu-X carboxypeptidase were shown to interact with ORF8, while the other glycoprotein GP36b with nsp7. These proteins are involved in immune regulation and other metabolic processes (22). These studies present likely evidence that mutations in these ORFs interacting druggable host proteins could perturb host-pathogen interactome, host immune responses or modulate therapeutic efficacy of anti-viral agents. The conspicuous distribution of co-occurring or exclusive mutations in the non-structural proteins; ORF1ab, ORF3a, ORF8 and N proteins of SARS-CoV-2 could be indicative of one or more of the following: a) Perturbations within SARS-CoV-2 proteome; b) Human-SARS-CoV-2 interactome or, c) Alteration of pharmacological profile of either known, or newly designed anti-viral candidates.

The current work highlights specific mutations in the functional genome of SARS-CoV-2 occurring worldwide and evolution of SARS-CoV-2 across different regions. While earlier reports have suggested the dominance of P314L and D614G subtype, we show additional dominance of other exclusive mutations in various countries like India, England Spain, and Brazil (Figure 1, 2). While the exact role of such mutations in conferring disease phenotype still needs to be deciphered, the proposed structural-mutation correlations for some of these mutations point toward their role in governing functional genome pathogenicity of SAR-CoV-2. The mutually dependent or exclusive role of these crucial mutations in global isolates is indicative of clonal geo-distribution, which may affect infectivity of SARS-CoV-2 in localized population. The identification of potential signature mutations in the evolving SARS-CoV-2 genome will aid the development of vaccines, drugs and diagnostics.

## Author Contributions

SEH and SAR designed this research; SAR, JS and HS performed research; SEH, SAR, JS and HS analysed data; and JS, SAR and HS wrote the paper. The authors declare no competing interest.

## Acknowledgements

SEH is a JC Bose National Fellow, Department of Science and Technology, Government of India and Robert Koch Fellow, Robert Koch Institute, Berlin. JS acknowledges fellowship support under Young Scientist and HS under Women Scientist Schemes of Dept. of Health Research, India. The authors acknowledge that BioInception and Envirozyme Biotech provided the genomic data analysis and discovery platform. We would like to thank Dr. Sergio M. Cuesta (Cancer Research UK Cambridge Institute, University of Cambridge) and other anonymous reviewers for their critical review of this manuscript.

